# Modulating Brain Rhythms of Pain using Transcranial Alternating Current Stimulation (tACS)? A Sham-controlled Study in Healthy Human Participants

**DOI:** 10.1101/2020.06.16.154112

**Authors:** Elisabeth S. May, Vanessa D. Hohn, Moritz M. Nickel, Laura Tiemann, Cristina Gil Ávila, Henrik Heitmann, Paul Sauseng, Markus Ploner

## Abstract

Pain protects the body. However, pain can also occur for longer periods without serving protective functions. Such chronic pain conditions are difficult to treat. Thus, a better understanding of the underlying neural mechanisms and new approaches for the treatment of pain are urgently needed. Here, we investigated a causal role of oscillatory brain activity for pain and explored the potential of transcranial alternating current stimulation (tACS) as a new treatment approach for pain. To this end, we investigated whether tACS can modulate pain and pain-related autonomic activity in 29 healthy human participants using a tonic heat pain paradigm as an experimental model of chronic pain. In 6 recording sessions, participants received tACS over prefrontal or somatosensory cortices at alpha or gamma frequencies or sham tACS. During tACS, pain ratings and autonomic responses were collected. TACS did not modulate pain intensity, the stability of pain ratings or the translation of the noxious stimulus into pain. Likewise, tACS did not change autonomic responses. Bayesian statistics further indicated a lack of tACS effects in most conditions. The only exception was alpha tACS over somatosensory cortex where evidence for tACS effects was inconclusive. Taken together, the present study did not find significant tACS effects on tonic experimental pain in healthy human participants. However, considering the conceptual plausibility of using tACS to modulate pain and the urgent need for novel pain treatments, further tACS studies are warranted. Based on the present findings, such studies might apply refined stimulation protocols targeting alpha oscillations in somatosensory cortices.

## 1. Introduction

Pain serves to protect the body. It signals threat and initiates behavioral adaptations and learning processes to limit and avoid harm. However, pain can also occur for extended periods of time without protecting the body. In such chronic pain conditions, pain represents a highly disabling disorder. With a prevalence of 20-30 %, chronic pain is a leading cause of disability worldwide and imposes an enormous burden on patients, health care systems, and society [14,36]. Current treatment approaches are often insufficient and can cause serious side effects as indicated by the Opioid crisis in the United States [31]. Moreover, the development of pain therapeutics is stagnating [40]. Thus, novel approaches for the treatment of chronic pain are urgently needed [19,34,40].

Recent insights into the brain mechanisms of pain open new perspectives for the development of novel pain treatments. Accumulating evidence indicates that pain is closely associated with neural oscillations at different frequencies and locations [32]. In particular, changes of neural oscillations at alpha (8 – 13 Hz) and gamma (30 – 100 Hz) frequencies in somatosensory and prefrontal brain areas have been related to longer-lasting experimental pain and chronic pain (e.g. [11,24,27,30,39,50]). Moreover, animal studies relying on optogenetics and invasive electrical stimulation have indicated that these changes of neural oscillations are causally involved in generating pain [43,51]. Thus, modulating neural oscillations to eventually modulate pain is a promising novel approach for the treatment of pain [15].

Transcranial alternating current stimulation (tACS) is an emerging neuromodulation technique which aims at non-invasively modulating neural oscillations in the human brain. During tACS, a weak alternating current is applied to the scalp with the goal of entraining neural oscillations at the stimulation frequency [33,49]. The appeal of the tACS is that it is non-invasive, safe, cost-efficient, and potentially mobile which allows for broad applications in clinical settings [15]. Thus, although the exact mechanisms and the effectiveness of tACS are still debated (Liu et al., 2018; Voroslakos et al., 2018), tACS is increasingly explored as a new treatment approach for neuropsychiatric disorders [19,42,44].

To date, two studies have investigated whether tACS can modulate pain [1,5]. Both studies employed tACS targeting somatosensory alpha oscillations. One of the studies assessed tACS effects on pain in patients suffering from chronic low back pain [1]. Overall, the results did not show tACS effects on pain. However, tACS induced increases in alpha oscillations, which correlated with changes in pain severity. The other study investigated tACS effects on brief experimental pain in healthy human participants [5]. Overall, the study did not show tACS effects on pain either. However, the findings indicated that tACS can reduce pain when expectations of upcoming pain are uncertain. The two studies thus provided preliminary evidence that tACS at alpha frequencies over somatosensory areas can potentially yield analgesic effects.

Gamma oscillations might represent another promising target for modulatory pain treatment approaches. Gamma oscillations in prefrontal brain areas encode pain intensity during tonic experimental and chronic pain [24,27,39]. In addition, gamma oscillations reliably track inter- and intraindividual variations of brief experimental pain in humans and rodents [16]. Furthermore, the optogenetic induction of gamma oscillations in the primary somatosensory cortex led to enhanced pain behavior indicating a causal role of gamma oscillations for pain [43]. However, although tACS at gamma frequencies is feasible [3,42], no study has explored the effects of gamma tACS on pain so far.

Here, we further explored the potential of tACS to modulate pain. To assess tACS effects on longer-lasting pain, we applied a tonic heat pain paradigm which resembles chronic pain conditions more closely than usual phasic pain stimuli [32]. Based on previous evidence on the role of neural oscillations in the processing of pain [1,5,24,27,30,35,39], we systematically applied tACS at alpha and gamma oscillations over somatosensory and prefrontal cortices in healthy human participants. To control for unspecific tACS effects, we additionally performed sham stimulation at both locations.

## 2. Materials and Methods

### 2.1 Participants

*A priori* sample size calculations using G*Power [13] determined a sample size of 28 participants for a repeated measures analysis of variance (RM ANOVA) design with 6 conditions (see below), a power of 0.95, an alpha of 0.05, and medium effect sizes of f = 0.25. This corresponds to an η^2^ (proportion variance explained) of 0.06 [9]. Based on these calculations, the final sample comprised 29 participants (all right-handed, 13 females, age: 25.7 ± 4.0 years [mean ± SD]). To this end, 39 healthy human participants were recruited. Ten participants were excluded during the course of experiment due to the absence of pain (n=3) or intolerable pain (n=3) during the first session, technical issues (n=2), or meeting exclusion criteria during one of the sessions (n=2).

Inclusion criteria were age above 18 years and right-handedness. Exclusion criteria were pregnancy, neurological or psychiatric diseases, severe internal diseases including diabetes, skin diseases, current or recurrent pain, regular intake of medication (aside from contraception, thyroidal and, in one case, antiallergic medication), previous surgeries at the head or spine, previous syncopes or head traumas resulting in unconsciousness or concussion, metal or electronic implants, and any previous side effects associated with thermal, electrical, or magnetic stimulation. None of the included participants showed signs of clinical anxiety or depression according to the Hospital Anxiety and Depression Scale (HADS) [52] with a cut-off of 8/21 [8] (anxiety: 2.5 ± 2.0 [mean ± SD]; depression: 0.7 ± 0.9 [mean ± SD]).

Prior to any experimental procedures, all participants gave written informed consent. The study protocol was approved by the local Ethics Committee of the Medical Faculty of the Technical University of Munich and pre-registered at ClinicalTrials.gov (NCT03805854). The study was conducted in accordance with the latest version of the Declaration of Helsinki and recent consensus guidelines for the application of tACS in humans [3].

### 2.2 Paradigm

In a within-subject design, each participant took part in 6 recording sessions. Sessions were separated by at least 24 hours and comprised a fixed sequence of events (Fig. 1A). In each session, tACS was applied over prefrontal or somatosensory cortices (Fig. 1B) using alpha frequency (10 Hz) stimulation, gamma frequency (80 Hz) stimulation, or sham stimulation (Fig. 1C). Concurrently, a tonic heat pain stimulus of varying intensity was applied to the left hand. During stimulation, participants continuously rated the currently perceived pain intensity. In addition, autonomic responses (skin conductance and electrocardiogram) were measured. Before and after the stimulation, 5 min of resting state EEG were recorded using the tACS electrodes.

**Fig. 1.**
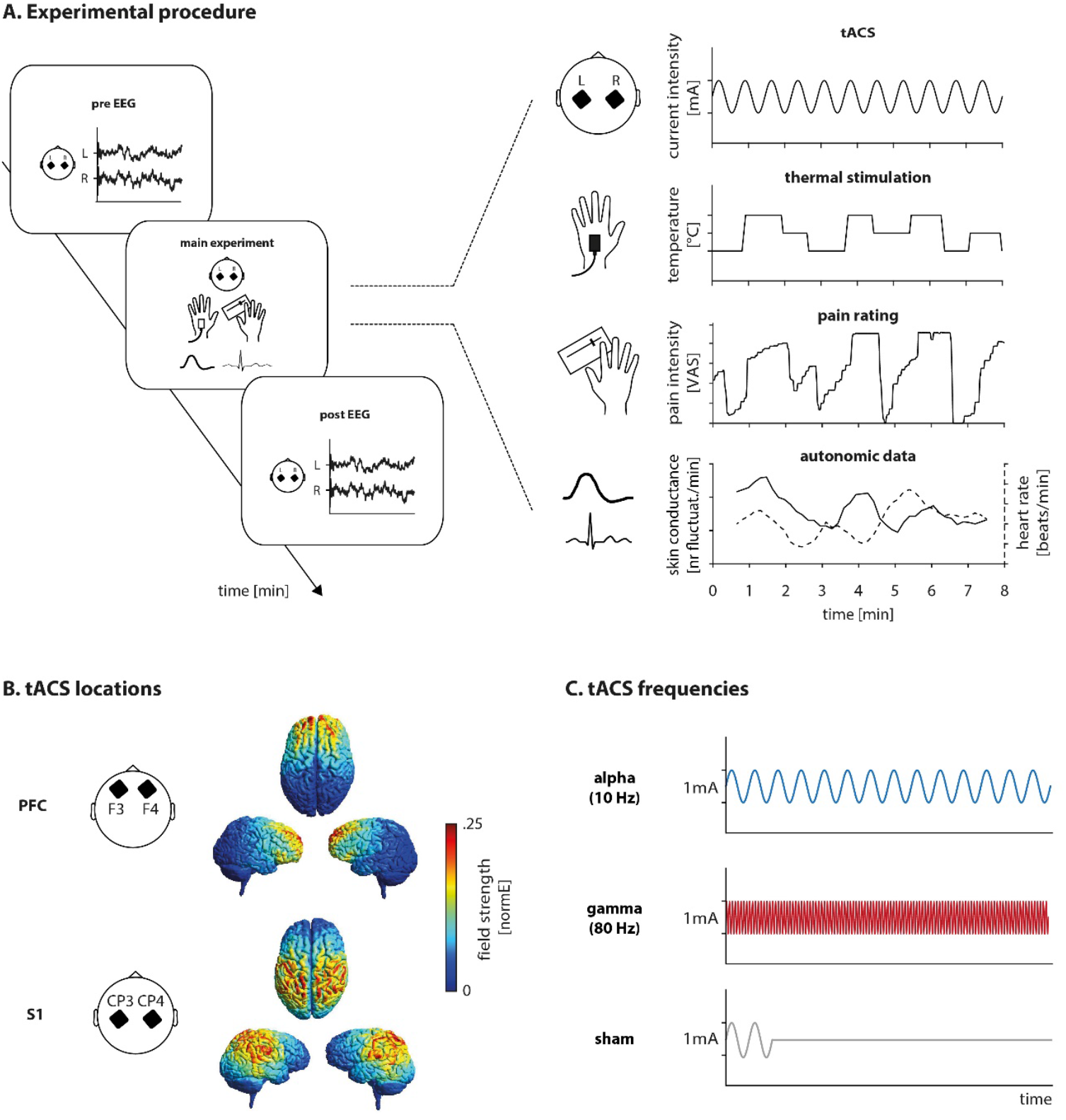
Paradigm. **(A)** Experimental Procedure. Each participant took part in 6 recording sessions which comprised a fixed sequence of events. During the main experiment, participants received transcranial alternating current stimulation (tACS) over prefrontal (PFC) or somatosensory (S1) cortices (B) using alpha, gamma, or sham stimulation (C) while a tonic heat pain stimulus of varying intensity was applied to the left hand. Concurrently, participants continuously rated the currently perceived pain intensity and autonomic responses (skin conductance and electrocardiogram) were measured. Before and after the main experiment, 5 min of resting state EEG were recorded using the tACS electrodes. **(B)** tACS locations. Using two 5*5 cm carbonized rubber electrodes placed according to the international 10-20 system, tACS of 1 mA intensity was applied over PFC (electrode positions F3 and F4) or S1 (electrode positions CP3 and CP4). Electrode placement was validated through simulations performed with SimNIBS 2.1 [38] using 1mA intensity, standard conductivity parameters, and the SimNIBS template head model. Simulations of the induced electrical field strength are shown on the right. **(C)** tACS frequencies. 1 mA tACS was applied at alpha or gamma frequencies or using sham stimulation. For alpha frequency stimulation, sinusoidal stimulation with a frequency of 10 Hz was applied. For gamma frequency stimulation, sinusoidal stimulation with 80 Hz frequency was applied. For sham stimulation, 30 s of 10 Hz sinusoidal stimulation were applied at the beginning of thermal stimulation only. All stimulations included 100 cycles fade-in and out. EEG, electroencephalography; L, left; PFC, prefrontal cortex; R, right; S1, primary somatosensory cortex; tACS, transcranial alternating current stimulation; VAS, visual analogue scale.

#### 2.2.1 Thermal stimulation

Tonic painful heat stimulation was applied to the participant’s left hand for 10 min using a thermode (TSA-II, Medoc, Ramat Yishai, Israel). Following an established paradigm [26–28], a predefined, fixed time course of stimulation (Fig. 1A) consisting of 9 plateaus with three temperature levels (low, medium, and high) was applied. Temperature levels were individually adjusted for each participant by adding 0.5, 0.8, or 1.1 °C to the individual pain threshold (see below), resulting in 3 intensity levels of thermal stimulation. The stimulation sequence consistent of 3 plateaus of 40, 50, and 60 s duration at each temperature level. The stimulation started from a baseline temperature of 40 °C and changed with a rate of 0.1 °C/s. All analyses were performed using an 8 min-time window beginning at the start of the first plateau.

Pain thresholds were determined for the left hand on the first recording day immediately before the pre-stimulation EEG resting state recording. Over the course of 3 min, participants continuously adjusted the thermode temperature to their individual pain threshold using two buttons of a computer mouse with their right hand. Depending on the side of the button press, the thermode temperature either increased or decreased with a rate of 0.5 C/s. The individual pain threshold was defined as the average stimulus intensity during the last 10 s and was used to determine individual temperature levels for all 6 recording days. Thus, temperature levels were individually adapted but constant across all conditions for each single participant. Mean pain threshold temperature was of 44.4 ± 1.7 °C [mean ± SD].

#### 2.2.2 Pain ratings

Simultaneously to thermal stimulation, participants continuously rated the currently perceived pain intensity on a visual analogue scale (VAS) ranging from 0 (“no pain”) to 100 (“worst tolerable pain”) using a custom-built finger span device with their right hand. The scale was simultaneously presented on a computer screen by a vertical orange bar, the height of which represented the current pain intensity. Pain ratings were sampled with a frequency of 1000 Hz by a BrainAmp ExG MR amplifier (Brain Products, Munich, Germany).

#### 2.2.3 Autonomic data

*Skin conductance* was recorded at the palmar distal phalanges of the left index and middle finger using Ag/AgCl electrodes connected to a GSR-MR module (Brain Products, Munich, Germany) with constant 0.5 V voltage. Participants were instructed not to move the hand during stimulation. Data were recorded in direct current (DC) mode with low-pass filtering at 250 Hz.

The *electrocardiogram* (ECG) was measured using a bipolar Ag/AgCl electrode montage with one electrode attached below the right clavicle and the other below the sternum. ECG data were band-pass filtered between 0.016 and 250 Hz. Both skin conductance and ECG were sampled at 1000 Hz using the BrainAmp ExG MR amplifier (Brain Products, Munich, Germany).

#### 2.2.4 Transcranial alternating current stimulation (tACS)

10 min of tACS was applied simultaneously to painful heat stimulation. tACS intensity was 1 mA for all participants and conditions. We employed a Neuroconn DC-STIMULATOR MR (Neuroconn, Ilmenau, Germany) and two carbonized rubber electrodes with a size of 5 x 5 cm. To validate electrode placement, electrical fields induced by a 1 mA transcranial current stimulation were simulated beforehand using SimNIBS 2.1 [38] with standard conductivity parameters and the SimNIBS template head model (Fig. 1B). For stimulation of the *prefrontal cortex (PFC),* electrodes were placed at positions F3 and F4 of the international 10-20 system. For stimulation of *somatosensory cortices (S1),* electrodes were attached at positions CP3 and CP4. Electrodes were fixed to the scalp using Ten20 conductive paste (D.O. Weaver, Aurora, CO, United States). Impedances were kept below 5 kΩ. For *alpha frequency* stimulation, a 10 min-sinusoidal stimulation with a frequency of 10 Hz was applied. For *gamma frequency* stimulation, 10 min-sinusoidal stimulation with 80 Hz frequency was applied. For *sham* stimulation, 30 s of 10 Hz sinusoidal stimulation were applied. All stimulations always included 100 cycles fade-in and -out. Fade-in always started with the beginning of thermal stimulation. Thus, during the 8 min-analysis window starting from the first plateau of thermal stimulation, participants received simultaneous, continuous tACS in the alpha and gamma frequency conditions, but no stimulation in the sham condition. For half of the participants, the 3 PFC sessions were performed first, followed by the 3 S1 sessions. For the other half, the order was reversed. Within the 3 sessions of each tACS location, the order of stimulation frequencies (alpha, gamma, sham) was counterbalanced.

#### 2.2.5 Pre- and post-stimulation EEG recordings

To quantify potential tACS effects on oscillatory brain activity outlasting the stimulation, 5 min of resting state brain activity were recorded immediately before and after tACS (pre- and post-EEG). Participants were asked to stay in a relaxed, wakeful state, without any particular task, keeping their eyes open and the gaze rested on a centrally presented fixation cross. EEG data were recorded using the same two electrodes used for tACS, i.e. placed at F3 and F4 for PFC sessions and at CP3 and CP4 for S1 sessions. Ag/AgCl electrodes attached to the nose and centrally on the forehead served as reference and ground, respectively. A bipolar Ag/AgCl electrode montage with electrodes below the outer canthus of the right eye and immediately below the hairline at the midline of the forehead was used to record eye movements. EEG data were sampled at 1000 Hz using the BrainAmp ExG MR amplifier (Brain Products, Munich, Germany) and bandpass-filtered between 0.016 and 250 Hz. Impedances were kept below 5 kΩ.

#### 2.2.6 Blinding

Due to the attachment of electrodes, participants and experimenters were not blinded with respect to the location of tACS. However, we aimed at a double-blind design with respect to the tACS frequency (alpha, gamma, sham). To this end, each session was conducted by a main experimenter who was unaware of the stimulation frequency and interacted with the participant and a second experimenter who operated the tACS device. At the end of each session, blinding of the participant was assessed using a short questionnaire consisting of three questions: (1) *“Did you have the impression that a **continuous** brain stimulation was applied today?”,* (2) *“Did you experience sensations at the scalp like tingling, prickling, or pulsing?”,* and (3) *“Did you experience light perceptions (phosphenes) like flickering?”.* Question 1 was answered using a forced-choice format *(yes/no),* whereas questions 2 and 3 were answered using a VAS ranging from 0 (“no”) to 10 (“very strongly”).

### 2.3 Data analysis

#### 2.3.1 tACS effects on pain

We first assessed whether tACS modulated pain perception (Fig. 2). For this purpose, the 8 min-pain rating and -temperature time courses were smoothed using a sliding-window approach with a window length of 1 s and a step size of 0.1 s. Smoothed pain rating and temperature time courses represented the basis of further analyses. Subsequently, analyses of tACS effects on *pain intensity* were performed. To investigate whether tACS influenced the overall pain level, we computed a summary measure of pain intensity by averaging pain ratings across the 8 min-interval and compared the resulting averages between conditions. To investigate whether tACS influenced pain intensity at any time during the 8 min of thermal stimulation, pain rating time courses were compared between alpha, gamma, and sham conditions in a time-resolved fashion. Lastly, we asked whether tACS might selectively alter pain intensity at certain temperature levels and compared the average pain intensity separately for low, medium, and high temperature levels.

**Fig. 2.**
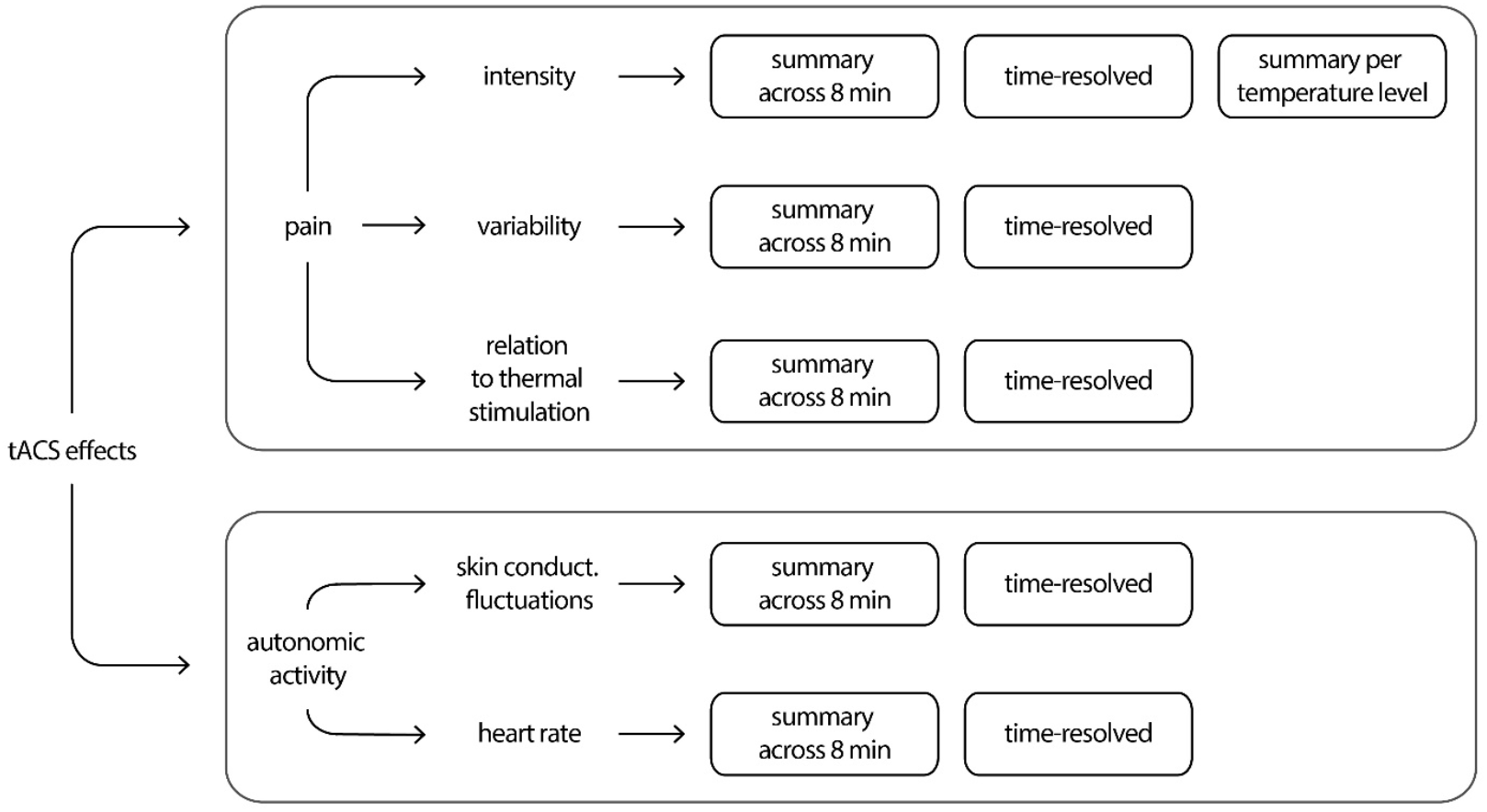
Analysis pipeline. The current study investigated effects of transcranial alternating current stimulation (tACS) on pain and pain-related autonomic activity. tACS effects on pain were investigated with respect to the intensity of pain, the variability of pain, and the relationship of pain to thermal stimulation. All variables were analysed across the entire 8 min analysis interval (summary measures), in a time-resolved fashion, as well as per temperature level in the case of pain intensity. tACS effects on autonomic activity were investigated using the number of skin conductance fluctuations (nSF) and the heart rate (HR). Again, variables were analyzed across the entire 8 min analysis window (summary measures) and in a time-resolved fashion. To detect tACS effects, all variables were compared between the three tACS conditions (alpha, gamma, sham) separately for both tACS locations (PFC, S1). PFC, prefrontal cortex; S1, primary somatosensory cortex.

Since tACS might also influence the stability of pain ratings or the translation of noxious stimuli into pain rather than pain directly, we next examined tACS effects on pain variability and the relation of pain to thermal stimulation. To this end, we first obtained summary measures across the 8 min-time course. For *pain variability,* the standard deviation of pain ratings was calculated across the entire time course. For the *relation of pain and thermal stimulation,* Pearson correlations between pain ratings and temperature were calculated. For additional time-resolved analyses, time courses of both measures were calculated by applying a sliding-window approach to the 8 min-pain rating and -temperature time courses using a window size of 60 s and a step size of 10 s [26]. Subsequently, the standard deviation of pain ratings and Pearson correlations between pain ratings and temperature were calculated for each window.

#### 2.3.2 tACS effects on autonomic activity

Next, we investigated whether tACS modulated pain-related responses of the autonomous nervous system, which are partially independent of pain perception [26]. Based on skin conductance and ECG recordings, the number of spontaneous skin conductance fluctuations (nSF) and the heart rate (HR) were computed and analyzed.

To obtain the nSF, skin conductance data were visually inspected for movement artifacts, low-pass filtered at 1 Hz using a fourth-order Butterworth filter, and downsampled to 500 Hz. Subsequently, a sliding-window approach with a window length of 60 s and a step size of 10 s was applied to the preprocessed skin conductance time series to obtain nSF time courses [26]. For every window, the nSF was determined by counting spontaneous fluctuations exceeding an amplitude criterion of 0.05 μS with respect to the proceeding trough. Windows contaminated with movement artifacts were discarded. To obtain the HR, ECG data were downsampled to 500 Hz and preprocessed using the Matlab toolbox PsPM, version 4.0.2 (bachlab.org/pspm). In PsPM, QRS complexes were detected and a continuous heart rate time series with a sampling frequency of 500 Hz was created by linearly interpolating the RR interval tachogram. Subsequently, the same sliding window approach used for skin conductance data was applied and the average HR was calculated for every window to obtain HR time courses [27].

First, tACS effects on the overall *strength of autonomic activity* were investigated by comparing summary measures obtained by averaging nSF and HR across the entire 8 min-time course between tACS conditions. Additionally, a time-resolved analysis was performed by comparing time courses of both measures across conditions.

#### 2.3.3 tACS effects on brain activity

We further investigated whether tACS induced neuronal changes outlasting the stimulation (offline effects). To this end, EEG data obtained before and after tACS were downsampled to 250 Hz. A visual artifact correction was performed, manually rejecting data segments contaminated by muscle activity. All analyses focused on the electrode contralateral to the stimulated hand, i.e. on F4 for the PFC and CP4 for the S1 electrode montage. In addition, the 60 s data segments closest to tACS were selected, i.e. the last min of the 5 min pre-stimulation EEG and the first min of the 5 min post-stimulation EEG. Data were cut into 1 s epochs with 50 % overlap and frequency specific power between 1 and 100 Hz was calculated using a Fast Fourier Transformation with a Hanning window resulting in a frequency resolution of 1 Hz. Subsequently, power spectra were averaged across all epochs for pre- and post-stimulation EEGs separately. Pre-stimulation power spectra were then subtracted from post-stimulation power spectra for each of the 6 conditions and each participant individually. During statistical analyses, these difference power spectra were compared between the active tACS conditions (PFC/S1 alpha/gamma stimulation) and the respective sham conditions (see below). As a control, the same analysis was performed calculating difference power spectra based upon the complete 5 min pre- and post-EEG data.

### 2.4 Statistical analyses

Statistical analyses were performed using Matlab (Mathworks, Natick, MA), the Matlab toolbox Fieldtrip [29], IBM SPSS Statistics for Windows (SPSS), version 26 (IBM Corp., Armonk, NY), and the statistical software package JASP, version 0.11.1 (JASP Team, 2019). Since Shapiro-Wilk tests indicated that some variables were not normally distributed, non-parametric tests were used for statistical analysis. These included Cochran’s Q-tests, Friedman tests, and nonparametric cluster-based permutation statistics [22,23]. Post hoc tests with Bonferroni correction were conducted when necessary and included McNemar tests for Cochran’s Q-tests and Wilcoxon matched-pairs signed rank tests for Friedman tests. Cluster-based permutation tests based on f tests were followed up by pairwise post hoc cluster-based permutation tests based on t-tests. Additionally, Bayesian RM ANOVAs were performed to complement analyses relying on null-hypothesis significance testing [7]. They were followed up by post hoc Bayesian dependent samples t-tests for analyses yielding moderate or strong evidence for either the null hypothesis of no tACS effect or the alternative hypothesis.

#### 2.4.1 Blinding

The blinding of participants was examined using a Cochran’s Q-test for question 1, which compared the frequency of yes responses across all 6 experimental conditions. VAS scores from question 2 and 3, which addressed the intensity of skin sensations and phosphenes, respectively, were investigated using Friedman tests. To investigate whether skin sensations and/or phosphenes differed between tACS locations, data from all frequency conditions were aggregated for each location and then compared using a Friedman test with the within-subjects factor location (PFC, S1). To investigate whether skin sensations and/or phosphenes differed between tACS frequencies, data from the three frequency conditions (alpha, gamma, sham) were compared using a Friedman test with the within-subjects factor frequency for both locations separately. All subsequent analyses were conducted separately for the PFC and S1 location as blinding questionnaires indicated that participants were successfully blinded for the S1 but not for the PFC condition.

#### 2.4.2 tACS effects on pain, autonomic activity, and brain activity

Friedman tests with the factor frequency (alpha, gamma, sham) were used to compare summary measures of pain intensity, pain variability, the relation of pain to thermal stimulation as well as nSF and HR. They were also used to investigate summary measures of pain intensity for each temperature level.

Cluster-based permutation statistics clustering across time were used to investigate tACS effects on time courses of pain intensity, pain variability, and the relation of pain to thermal stimulation, as well as time courses of nSF and HR in a time-resolved fashion. Specifically, time courses were compared between alpha, gamma, and sham conditions using cluster-based permutation statistics based on dependent samples f-tests, clustering across time.

Cluster-based permutation statistics clustering across frequencies were used to investigate tACS offline effects on brain activity power spectra. To this end, pre- and post-stimulation difference power spectra from the electrode contralateral to thermal stimulation were compared between the active tACS conditions and the respective sham conditions using nonparametric cluster-based permutation statistics based on dependent samples t-tests, clustering effects across frequencies. Specifically, PFC alpha and gamma conditions were compared to the PFC sham condition and S1 alpha and gamma conditions were compared to the S1 sham condition resulting in 4 pairwise comparisons. When investigating the effect of alpha frequency stimulation, this analysis was applied to a frequency band from 8 to 12 Hz. When investigating gamma frequency stimulation, the frequency band was 70 to 90 Hz.

To *control for multiple comparisons*, all p values were subjected to false discovery rate (FDR) control of Type I error [6]. Corrections were conducted separately for pain, autonomic activity, and brain activity considering the number of all statistical analyses performed for the respective measure (see Fig. 2 for an overview for pain and autonomic activity). This resulted in an FDR control for 18 statistical tests (9 tests x 2 tACS locations) for pain ratings, an FDR control for 8 statistical tests (2 tests x 2 tACS locations x 2 autonomic measures) for autonomic activity, and an FDR control for 4 statistical tests (2 tests x 2 tACS locations) for brain activity. Throughout the manuscript, corrected p-values are reported. Uncorrected p-values for all analyses are summarized in Supplementary Table S1. If not stated otherwise, statistical tests were performed two-sided with a significance level (α) of p < .05.

Analyses relying on null-hypothesis significance testing were complemented by *Bayesian RM ANOVAs.* The Bayesian approach to hypothesis testing considers the likelihood of the observed data under the null and the alternative hypothesis. The comparison of the resulting probabilities is reflected by the Bayes Factor (BF01 = likelihood of the data given the H0 / likelihood of the data given the H1) [7,17]. Thus, Bayes factors allow to specifically evaluate evidence in favor of the null hypothesis. Bayesian RM ANOVAs were performed for the pain intensity summary measure as well as the average nSF and HR across 8 min. As before, the analyses included the factor frequency (alpha, gamma, sham) and were conducted separately for both tACS locations. For all effects, JASP default prior options were chosen.

### 2.5 Subgroup analyses

To further investigate potential tACS effects for those participants presumably responding best to the brain stimulation, we repeated pain and autonomic activity analyses for two subgroups. In a *responder analysis,* analyses were performed for a subgroup of participants showing the strongest evidence for frequency specific tACS offline effects on brain activity quantified by calculating individual alpha and gamma responder ratios. These were based on EEG data from the last min pre-stimulation and the first min post-stimulation. To this end, EEG data from the electrode contralateral to stimulation were cut into 1 s epochs with 50 % overlap, power spectra were calculated using a Fast Fourier Transformation and a Hanning window and averaged across all epochs. To investigate alpha frequency stimulation effects, power values were averaged between 8 and 12 Hz. To investigate gamma frequency effects, power values were averaged between 70 and 90 Hz. Adopting the approach of D’Atri et al. [10], the ratio between post- and pre-EEG power values was then calculated for the active tACS conditions (alpha, gamma) and normalized by the ratio derived for the respective sham condition:

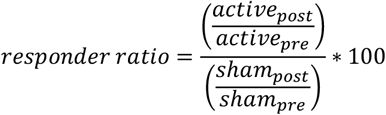

These calculations were performed separately for the alpha and gamma conditions and the PFC and S1 electrodes, resulting in four responder ratios per participant. Participants with responder ratios above 100 were classified as responders [10]. Overall, 15 of 29 participants were classified as PFC alpha responders, 18 as PFC gamma responders, 18 as S1 alpha responders, and 13 as S1 gamma responders, resulting in four condition- and frequency-specific subgroups.

In a *peak frequency informed analysis*, a selection was made based on the proximity of the stimulation frequency to the frequency of endogenous oscillations in the stimulated brain area, since this proximity can influence the degree of neural entrainment induced by tACS [49]. As power spectra displayed clear peaks of endogenous oscillations at alpha frequencies over S1 exclusively, this analysis was only applied to the S1 alpha condition. Specifically, analyses were performed for a subgroup of participants whose individual alpha peak frequency (IAF) was closest to the 10 Hz-stimulation frequency. To estimate IAFs, the entire 5 min, artifact-cleaned pre-stimulation EEG from the S1 alpha condition was cut into 10 s epochs with 50 % overlap, calculating power spectra using Fast Fourier Transformation and Hanning windows. A longer epoch length of 10 s was chosen to increase the frequency resolution to 0.1 Hz. Based on the averaged power spectra, the IAF was defined for each participant as the local power maximum in the frequency range between 8 and 13 Hz. Subsequently, peaks were visually controlled and corrected whenever reasonable. Following this procedure, IAF peaks could be identified for 26 of 29 participants. Next, the 50 % of participants whose IAF was closest to the stimulation frequency of 10 Hz were selected, resulting in a subgroup of 13 participants.

Pain and autonomic activity analyses outlined above were repeated for the two subgroups with slight adjustments. First, three-way comparisons were replaced by two-way comparisons (e.g. S1 alpha vs. S1 sham instead of S1 alpha vs. S1 gamma vs. S1 sham) because the number of responders differed between stimulation conditions (e.g. responders_S1alpha_ ≠ responder_S1gamma_) and IAFs were determined for the S1 alpha condition only. Thus, cluster-based permutation statistics were based on dependent sample t-tests instead of f-tests and the within-subjects factor stimulation was reduced from 3 to 2 levels for all tests. Second, for the peak frequency informed analysis, analyses were repeated for the S1 alpha frequency condition only, contrasting effects in the S1 alpha frequency condition to those obtained in the S1 sham condition. Third, the applied FDR correction was adjusted to the number of statistical tests conducted. For pain ratings, this resulted in an FDR control for 36 tests (14 tests x 2 tACS locations) and for 9 tests (9 tests x 1 tACS locations) in the responder and peak frequency informed analysis, respectively. For autonomic activity, an FDR control for 16 tests (4 tests x 2 tACS locations x 2 autonomic measures) and for 4 tests (4 tests x 1 tACS location) was applied.

### 2.6 Data availability

All data and scripts will be available upon final publication.

## 3. Results

To investigate whether tACS can modulate pain, participants took part in 6 recording sessions. During each session, tACS was applied at one of two locations (over PFC or S1) and at one of three frequencies (alpha [10 Hz], gamma [80 Hz], sham). Concurrently, a tonic heat pain stimulus of varying intensity was applied to the left hand.

### 3.1 Participants were blinded for tACS over S1, but not over PFC

After each session, the blinding of participants was assessed using questionnaires (Supplementary Fig. S1). When asked whether a continuous stimulation was applied or not, participants’ reports did not differ between tACS conditions (χ^2^(5) = 7.46, p = .189). Likewise, skin sensations did neither differ between tACS locations (χ^2^(1) = .75, p = .385) nor between tACS frequencies for either of the locations (PFC: χ^2^(2) = 2.11, p = .348; S1: χ^2^(2) = 0.75, p = .688). However, phosphenes were stronger for tACS over PFC than over S1 (χ^2^(1) = 7.23, p = .007). In addition, phosphenes differed between frequencies for tACS over PFC, but not over S1 (PFC: χ^2^(2) = 8.90, p = .012; S1: χ^2^(2) = 0.19, p = .910). Post hoc tests showed that phosphenes were significantly stronger in the PFC alpha condition than in the PFC gamma condition (Z = −3.13, p = .006). Hence, participants were successfully blinded for tACS over S1 but not for tACS over PFC. Thus, all further analyses investigated tACS effects separately for PFC and S1 locations.

### 3.2 tACS did not modulate pain

We first investigated whether tACS influenced pain intensity averaged across the entire 8 min of thermal stimulation. To this end, we compared average pain intensity during alpha, gamma, and sham tACS for both locations (Fig. 3). The results did not show any statistically significant differences, neither during tACS over PFC nor during tACS over S1 (p > .05 for all tests; see Supplementary Table S1 for test statistics and uncorrected p-values of all pain analyses). We further assessed whether tACS influenced pain intensity at any time during the 8 min of thermal stimulation. To this end, we compared pain intensity time courses during alpha, gamma, and sham stimulation for both tACS locations (Fig. 3). For both PFC and S1, cluster-based permutation tests did not show significant differences in pain intensity at any time (p > .05 for all clusters). We further asked whether tACS might selectively alter pain intensity at certain temperature levels. For instance, tACS might particularly modulate pain at the lowest level at which pain ratings are closest to pain threshold and possibly most uncertain. We therefore compared the average pain intensity separately for low, medium, and high temperature levels (Supplementary Fig. S2). However, no significant tACS effects on pain intensity were found for any temperature level (p > .05 for all tests). Lastly, we asked whether tACS might influence the stability of pain ratings or the translation of noxious stimuli into pain rather than pain intensity directly. To this end, we investigated whether tACS influenced the variability of pain or the relation of pain to thermal stimulation (Fig. 4). Comparisons of summary measures across 8 min did not yield significant tACS effects on pain variability or the relationship of pain to thermal stimulation (p > .05 for all tests). Likewise, time-resolved analyses of both measures did not show any significant tACS effects at any time (p > .05 for all clusters). Taken together, we did not find tACS effects on different measures of tonic pain.

**Fig. 3.**
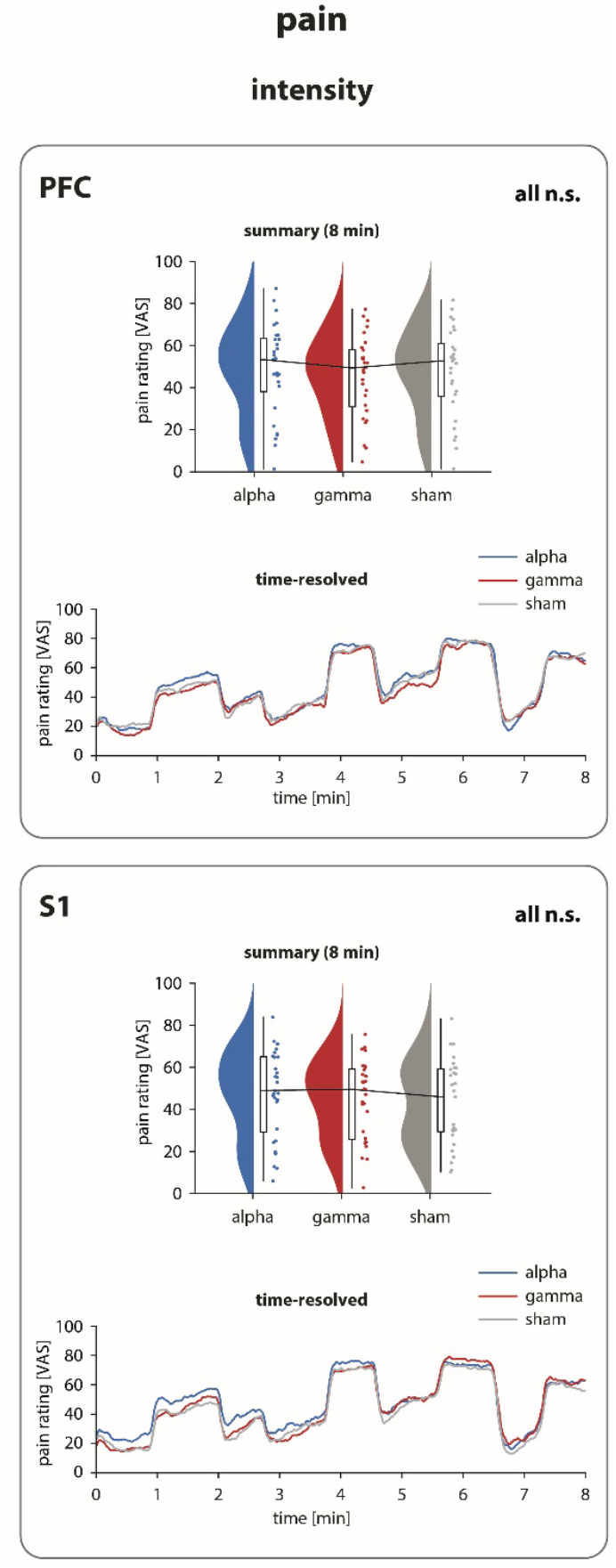
tACS effects on pain intensity. tACS effects on pain intensity are shown separately for PFC (upper panel) and S1 (lower panel) tACS locations. For both locations, upper rows display summary measures obtained by averaging pain ratings (0-100; VAS) across the 8 min analysis window. Raincloud plots [2] show un-mirrored violin plots displaying the probability density function of the data, boxplots, and individual data points. Boxplots depict the sample median as well as first (Q1) and third quartiles (Q3). Whiskers extend from Q1 to the smallest value within Q1 – 1.5* interquartile range (IQR) and from Q3 to the largest values within Q3 + 1.5* IQR. Lower rows depict time-resolved analyses of pain rating time courses in the alpha, gamma, and sham tACS conditions. None of the analyses revealed significant differences between alpha, gamma, and sham stimulation indicating no tACS effects on the perceived pain intensity (N = 29; PFC_summary_: p = .885, PFC_time-resolved_: no cluster found, S1_summary_: p = .864, S1_time-resolved_: p = .857; Friedman tests and cluster-based permutation statistics; FDR-corrected p-values). n.s., not significant; PFC, prefrontal cortex; S1, somatosensory cortex; tACS, transcranial alternating current stimulation; VAS, visual analogue scale.

**Fig. 4.**
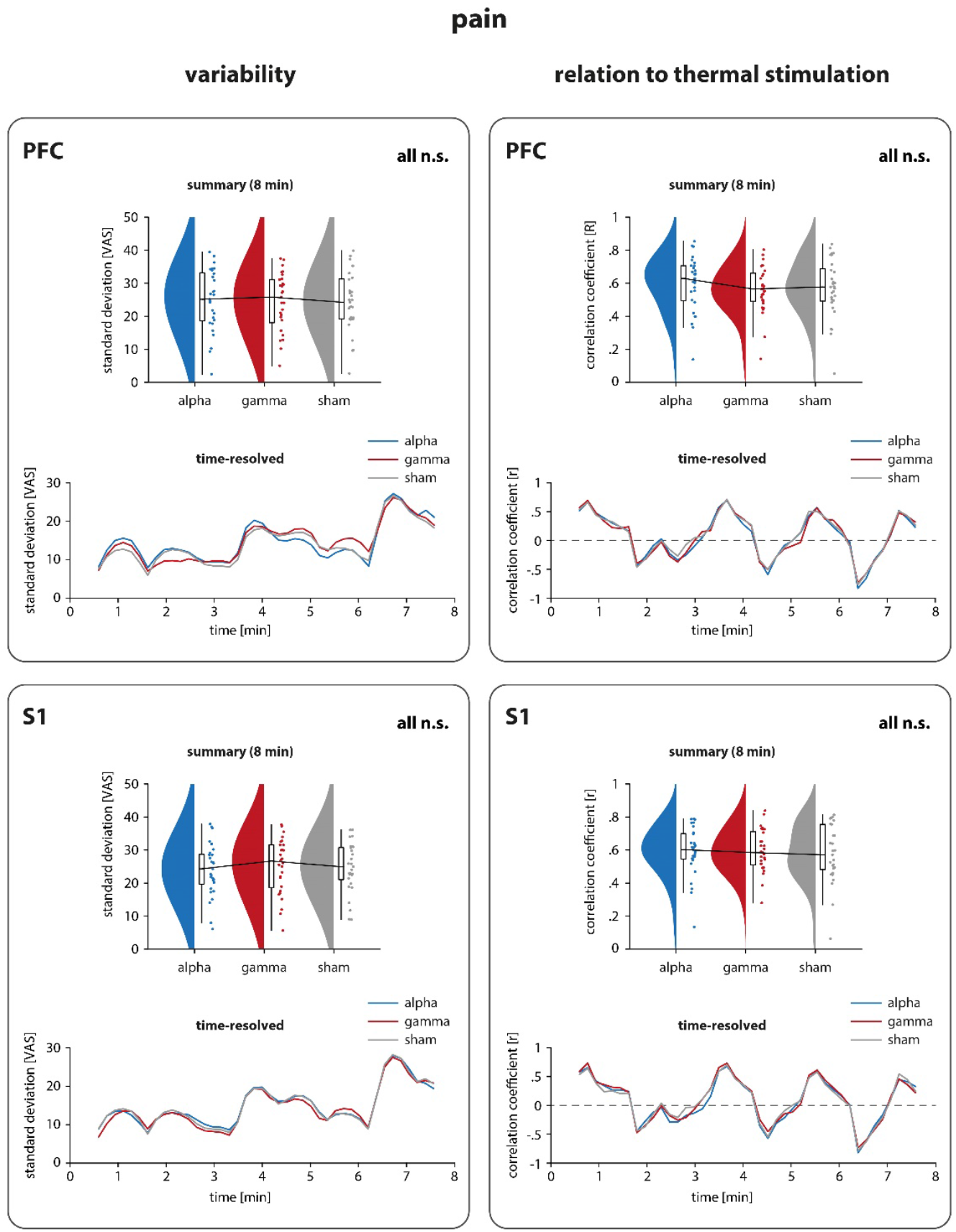
tACS effects on pain variability and the relation of pain to thermal stimulation. In line with Fig. 3, tACS effects on pain variability and the relation of pain to thermal stimulation are shown separately for PFC and S1 tACS locations. Results of the analyses based on summary measures obtained across the 8 min analysis window as well as results of the time-resolved analyses are shown. None of the analyses revealed significant differences between alpha, gamma, and sham stimulation, indicating no tACS effects on the stability of pain ratings or the translation of the noxious stimulus into pain (N = 29; pain variability: PFC_summary_: p = .864, PFC_time-resoked_: p = .857, S1_summary_: p = .857, S1_time-resolved_: p = .864; relation of pain to thermal stimulation: PFC_summary_: p = .943, PFC_time-resolved_: p = .857, S1_summary_: p = .943, S1_time-resolved_: p = .857; Friedman tests and cluster-based permutation statistics; FDR-corrected p-values). n.s., not significant; PFC, prefrontal cortex; S1, somatosensory cortex; tACS, transcranial alternating current stimulation; VAS, visual analogue scale.

Next, we used Bayesian statistics to evaluate direct evidence for a lack of tACS effects on pain intensity. We specifically performed Bayesian RM ANOVAs with the factor tACS frequency (alpha, gamma, sham) for both PFC and S1 locations. These analyses resulted in a Bayes factor (BF_01_) which quantifies the relative likelihood of the data given the null hypothesis of no tACS effect over the alternative hypothesis postulating tACS effects. BF_01_ values below 0.33 are commonly classified as evidence for the alternative hypothesis, values from 0.33 to 3 are classified as inconclusive evidence, and values above 3 are classified as evidence for the null hypothesis [18]. The analysis of pain intensity resulted in a BF_01_ of 5.727 for the PFC and a BF_01_ of 2.082 for the S1 location, indicating that evidence for the null hypothesis was moderate for the PFC but inconclusive for the S1 location. To follow up the inconclusive result for tACS over S1, we performed pairwise comparisons between tACS frequencies (alpha, gamma, sham) using Bayesian dependent samples t-tests. These revealed moderate evidence for the null hypothesis when comparing the gamma and sham conditions (BF_01 gamma ≠ sham_ = 3.893) but inconclusive evidence for both comparisons entailing the alpha condition (BF_01 alpha ≠ sham_ = 0.935; BF_01 alpha ≠ gamma_ = 2.491).

Taken together, frequentist statistical analyses did not provide evidence for a modulation of tonic pain by tACS at alpha or gamma frequencies over prefrontal or somatosensory cortices. Bayesian analyses provided moderate evidence for a lack of tACS effects on tonic pain except for alpha tACS over somatosensory areas where evidence was inconclusive.

### 3.3 tACS did not modulate pain-related autonomic activity

We further examined whether tACS influenced pain-related activity of the autonomic nervous system. Such autonomic responses to noxious stimuli are partially independent of pain perception [26]. Moreover, autonomic responses are mediated by different brain mechanisms than pain perception [45]. To this end, we analyzed tACS effects on nSF and HR (Fig. 5). We did not find any tACS effects on these autonomic measures neither when summary measures across 8 min, nor when time courses were compared between tACS conditions (Fig. 5, p > .05 for all analyses; see Supplementary Table S1 for all test statistics and uncorrected p-values). In addition, we applied Bayesian RM ANOVAs to evaluate direct evidence for a lack of tACS effects on nSF and HR. Analyses of both measures provided moderate evidence for the null hypothesis for both the PFC and the S1 location (nSF: BF_01 PFC_ = 3.88, BF_01 S1_ = 4.36; HR: BF_01_ = 6.59, BF_01_ = 8.95).

**Fig. 5.**
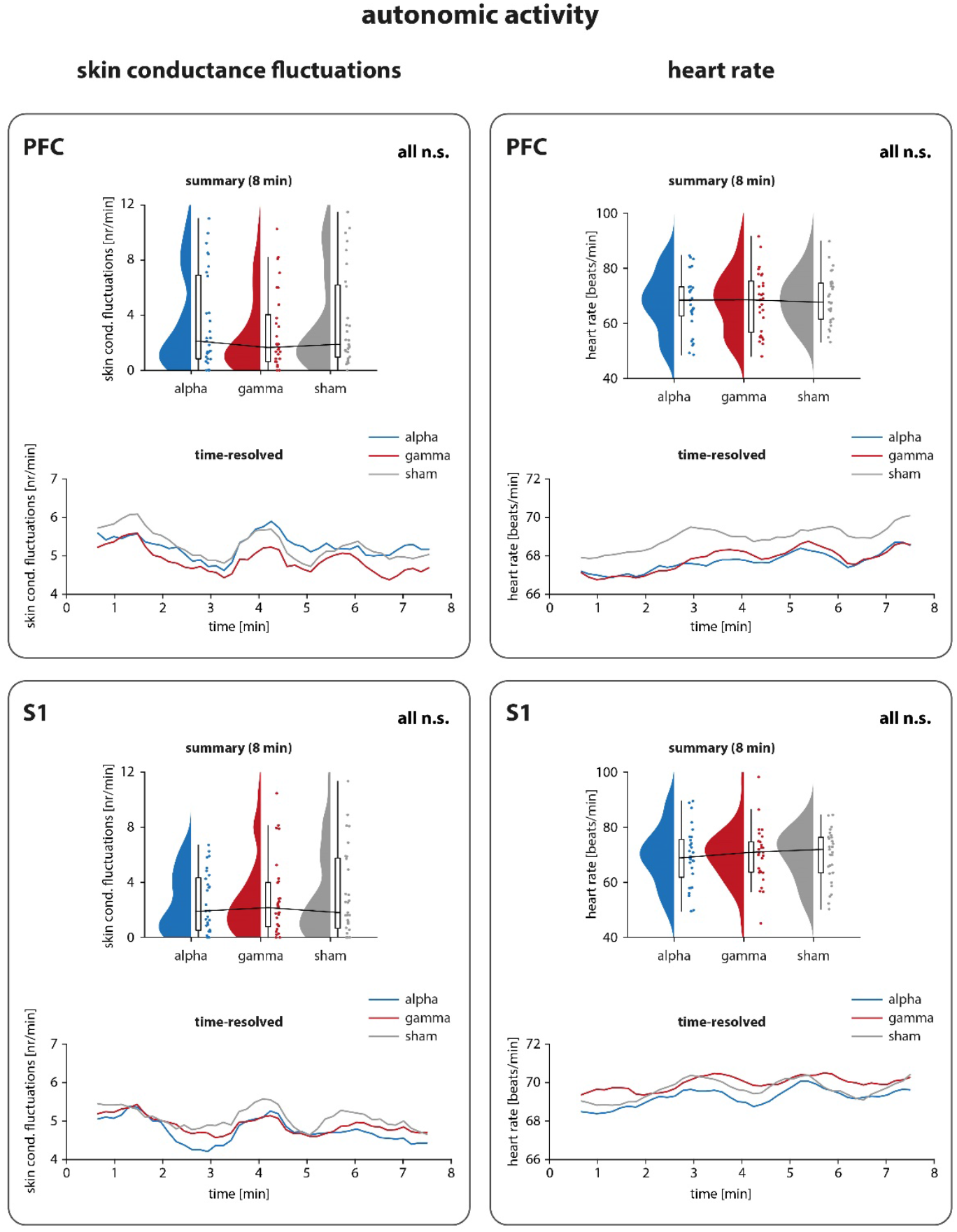
tACS effects on autonomic activity. In line with Fig. 3, tACS effects on the number of skin conductance fluctuations and the heart rate are shown separately for PFC and S1 locations. Results of the analyses based on summary measures obtained across the 8 min analysis window as well as results of the time-resolved analyses are shown. Due to the chosen windowing approach with a window length of 1 min and a step size of 10 s, time-courses of autonomic data span 7 min only and are depicted from 0.5 to 7.5 min. None of the analyses revealed significant differences between alpha, gamma, and sham stimulation, indicating no tACS effects on pain-related autonomic activity (N = 29; number of skin conductance fluctuations: PFC_summary_: p = 1.0, PFC_time-resolved_: no cluster found, S1_summary_: p = 1.0, S1_time-resolved_: no cluster found; heart rate: PFC_summary_: p = 1.0, PFC_time-resolved_: no cluster found, S1_summary_: p = 1.0, S1_time-resolved_: no cluster found; Friedman tests and cluster-based permutation statistics; FDR-corrected p-values). n.s., not significant; PFC, prefrontal cortex; S1, somatosensory cortex; tACS, transcranial alternating current stimulation.

Taken together, we did not find significant tACS effects on autonomic activity during tonic pain. Bayesian analyses provided moderate evidence for a lack of tACS effects on pain-related activity of the autonomic nervous system.

### 3.4 tACS did not yield outlasting effects on brain activity

The rationale of tACS is to modulate neural oscillations during tACS. Demonstrating such modulations directly requires the simultaneous measurement of neural oscillations during tACS. However, such online EEG measurements are heavily contaminated by tACS artifacts and their significance is therefore uncertain [49]. Apart from effects during tACS, some studies indicated that tACS can also yield effects on neural oscillations which outlast tACS [49]. To check for such offline effects as a potential indicator of the neural efficacy of tACS, we next investigated whether tACS induced neuronal changes outlasting the stimulation. To this end, we calculated power spectra of EEG activity during the last minute before and the first minute after stimulation (Fig. 6). We further calculated post – pre difference power spectra of the electrode contralateral to the thermal stimulation and compared them between the active tACS conditions (PFC/S1 alpha/gamma stimulation) and the respective sham conditions. Cluster-based permutation statistics did not show any significant clusters for tACS over PFC or S1 in the targeted frequency bands (p > .05 for all clusters, one-sided). Control analyses including the entire 5 min pre- and post-EEG data for power spectra calculation confirmed this finding (p > .05 for all clusters, one-sided). Hence, tACS did not evoke effects on brain activity outlasting the stimulation.

**Fig. 6.**
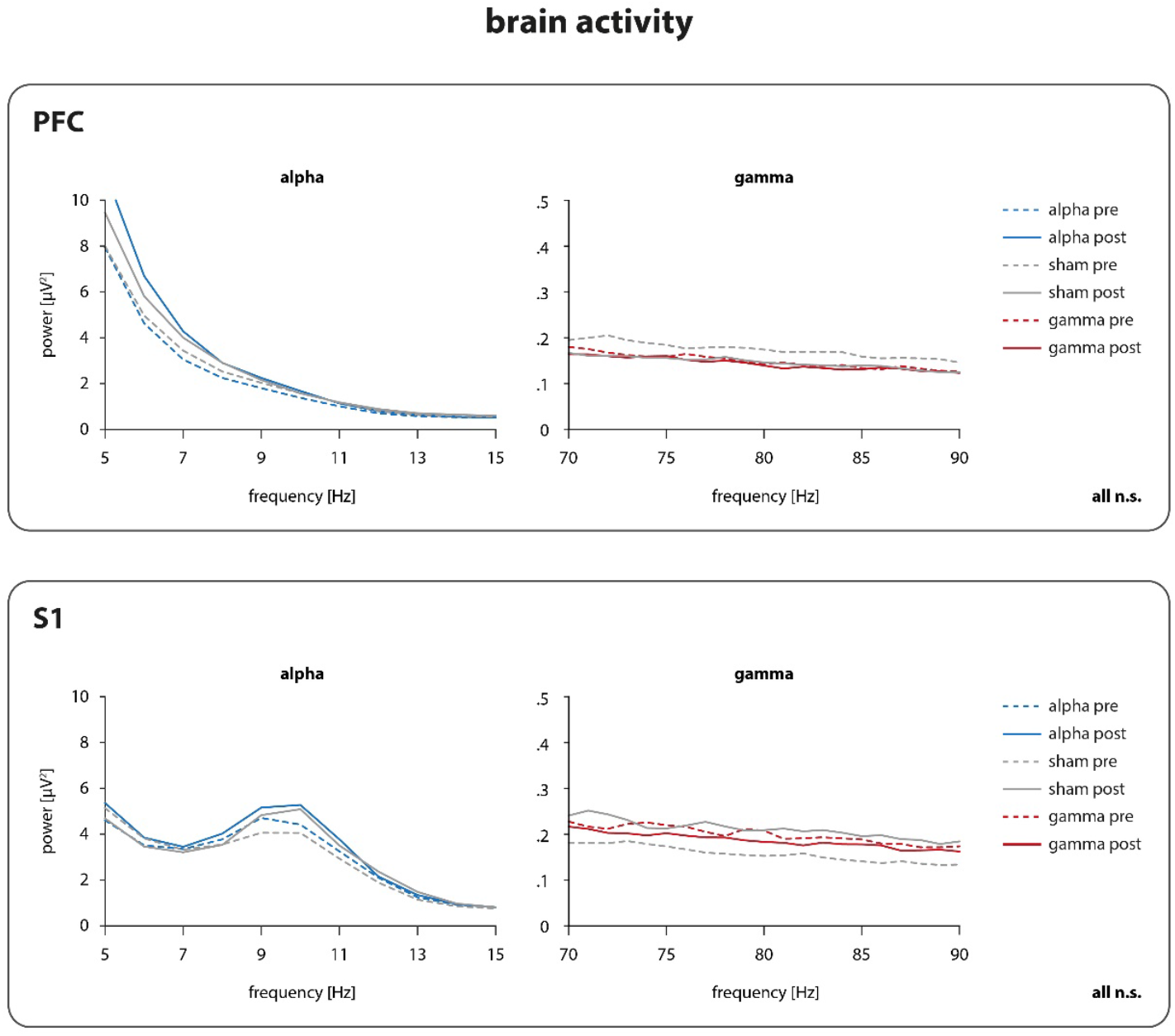
tACS effects on brain activity. Power spectra of pre- (dashed lined; based on last min of the recording) and post-stimulation EEGs (solid line; based on first min of the recording) are shown separately for PFC (upper panel) and S1 (lower panel) locations. Left and right plots display power spectra for alpha and gamma frequency bands, respectively. For statistical analysis, pre-stimulation power spectra were subtracted from post-stimulation power spectra for each of the 6 conditions and each participant individually (not shown here). Subsequently, the resulting difference power spectra were compared between the active tACS conditions and the respective sham condttbns (pFC_alpha_ vs PFC_sham_, PFC_gamma_ vs PFC_sham_, S1_alpha_ vs S1_sham_, S1_gamma_ vs S1_sham_). AnalySeS did not reveal significant power increases in the targeted frequency bands indicating that tACS did not evoke effects on brain activity outlasting the stimulation (N = 29; PFC_alpha_: no cluster found, PFC_gamma_: no cluster found, S1_alpha_: p = .240, S1_gamma_: no cluster found; Cluster-based permutation statistics; one-sided FDR-corrected p-values). EEG, electroencephalography; n.s., not significant; PFC, prefrontal cortex; S1, somatosensory cortex, tACS, transcranial alternating current stimulation.

### 3.5 tA CS did not influence pain and autonomic activity in subgroups of participants selected based on the responsiveness to tACS and individual peak alpha frequencies

Finally, we investigated whether tACS influenced pain or autonomic activity in selected participants who might have responded particularly strongly to tACS. We therefore performed two subgroup analyses. First, although the group-level analysis did not show tACS effects on brain activity, we tested for tACS effects in those participants who showed the highest frequency-specific post/pre ratio of brain activity *(responders).* Repeating the whole-group analyses for these subgroups did not reveal any tACS effects on pain or autonomic activity (p > .05 for all analyses). Second, we investigated tACS effects in those 50 % of participants whose individual alpha peak frequency was closest to the 10 Hz tACS alpha stimulation. As the individual alpha peak frequency can be reliably identified over S1 only, this analysis was only performed for the S1 alpha and S1 sham condition. Repeating the whole group analyses for these subgroups did not reveal any tACS effects on tonic pain or autonomic activity (p > .05 for all analyses).

## 4. Discussion

The current study systematically explored whether tACS can modulate pain and pain-related autonomic activity in healthy human participants using a tonic heat pain paradigm. In 6 recording sessions, participants received tACS over PFC or S1 using alpha, gamma, or sham stimulation while pain ratings and autonomic responses were collected. Analyses showed that tACS did not modulate the perceived pain intensity, the stability of pain ratings or the translation of the noxious stimulus into pain. Likewise, tACS did not influence autonomic responses. Bayesian statistics further supported a lack of tACS effects in most conditions including prefrontal and gamma tACS. The only exception was alpha tACS over S1 where evidence for tACS effects on tonic pain intensity was inconclusive. Thus, somatosensory alpha oscillations remain the most promising target for a modulation of tonic pain using tACS.

The present study complements two previous tACS studies in the context of pain [1,5]. Both studies applied 10 Hz alpha stimulation targeting somatosensory areas, which has also been done in the current study. One study indicated an analgesic effect of tACS on pain induced by brief experimental stimuli but only when pain intensity was uncertain [5]. The other study applied tACS to chronic back pain patients. The results did not show a tACS effect on overall pain levels. However, tACS induced changes of alpha activity which correlated with pain intensity [1]. In addition, exploratory analyses of autonomic activity recorded in this study indicated that heart rate variability was increased [35]. Our study extends these findings in two important aspects. First, we not only targeted somatosensory alpha oscillations but systematically assessed tACS effects at alpha and gamma frequencies over somatosensory and prefrontal brain areas. Second, to particularly explore tACS effects on longer-lasting pain, we employed a tonic heat pain paradigm. In addition, our study followed a careful design with strong methodological rigor. Sample size was based on *a priori* sample size calculation. Targets and frequencies of tACS were clearly motivated by previous studies on the role of neural oscillations in the processing of pain (May et al., 2018; Nickel, May, Tiemann, Schmidt, et al., 2017; Peng, Hu, Zhang, & Hu, 2014; Schulz et al., 2015). Sham conditions controlled for unspecific tACS effects. After each session, blinding was assessed using post-stimulation questionnaires. Extensive analyses of pain and autonomic activity were performed and adequately corrected for multiple comparisons. Finally, analyses were complemented by Bayesian statistics to strengthen the interpretability of negative findings [7]. However, complementing the weak tACS effects on pain in previous studies [1,5,35], we did not find significant tACS effects on tonic experimental pain in healthy human participants.

The lack of tACS effects on pain might be due to different factors. First, tACS might in general not be able to modulate neural oscillations in the brain. Although behavioral and, to a lesser extent, neural evidence for the effectiveness of tACS is growing [49], it is still debated whether currents applied in human tACS studies are sufficiently strong to pass through the skull and modulate brain activity [21,47]. Thus, we cannot rule out that the lack of tACS effects in the present study reflects a general lack of tACS effectiveness. Second, tACS might be able to modulate neural oscillations but neural oscillations might not be causally involved in the processing of pain. This seems unlikely since optogenetic and invasive electrical modulations of neural oscillations in somatosensory and prefrontal brain areas effectively modulate acute and chronic pain in animals [43,51]. Third, tACS might in principle be able to modulate neural oscillations and pain but the tACS parameters used in the present study were not optimal. To ensure that certain tACS paradigms are able to modulate neural oscillations, simultaneous EEG recordings are desirable. However, such simultaneous EEG recordings are heavily contaminated by tACS-induced artifacts [49]. In the present study, we therefore performed EEG recordings immediately after tACS. The results did not show significant changes of EEG activity after tACS. However, as EEG after-effects are not consistently observed [25,48], their lack does not preclude tACS effects during stimulation. It is nevertheless possible that the tACS parameters of the present study were not optimal for modulating pain and future studies might apply other, individualized parameters. Fourth, tACS might in principle be able to modulate neural oscillations and pain but with a different pain paradigm than the one chosen here. The tonic pain stimulus we applied evokes strong decreases in alpha oscillations in early somatosensory areas [27,39], which, together with previous tACS studies, led us to target S1 alpha oscillations using tACS. However, neuronal entrainment by tACS depends on the amplitude of the targeted oscillations [37,41]. Consequently, tasks which strongly suppress oscillations might be less susceptible for modulations using tACS. Future studies might therefore use other pain paradigms which yield weaker and/or shorter suppressions of oscillations.

In conclusion, our findings do not provide evidence that tACS can modulate tonic experimental pain in healthy human participants. This might indicate a fundamental inability of tACS to reach the brain and/or modulate pain. Alternatively, it might point out the need to optimize the experimental setup in further studies. For instance, we chose standardized tACS locations and stimulation intensities with the goal of a broad clinical usability in mind. However, individualized tACS parameters might be more effective. Future studies might therefore consider the participant’s individual anatomy as well as the individual peak frequency of the targeted neural oscillations [12,20]. In addition, increasing tACS intensity, duration, and/or perform repeated stimulation sessions might increase its neural efficacy. With respect to gamma oscillations, transcranial random noise stimulation (tRNS) [4] rather than sinusoidal stimulation at a specific frequency might be more effective due to the broad-band, burst-like nature of neuronal gamma activity [46]. Considering the urgent need for novel pain treatments, the conceptual plausibility and potentially broad clinical applicability of tACS to modulate pain, and the relative lack of alternatives, we feel that such follow-up studies are warranted. However, just like any single chronic pain treatment approach, tACS will likely not represent a stand-alone treatment but rather a valuable part of a combined bio-psycho-social treatment approach. Based on the present and previous studies [1,5,35], the most promising target for future tACS studies are alpha oscillations in somatosensory cortices.

## Acknowledgments

The study was supported by the Deutsche Forschungsgemeinschaft (PL 321/10-2, PL 321/11-2) and the Studienstiftung des deutschen Volkes.

The authors have no conflict of interest to declare.

## Supplementary Materials

**Supplementary Fig. 1.**
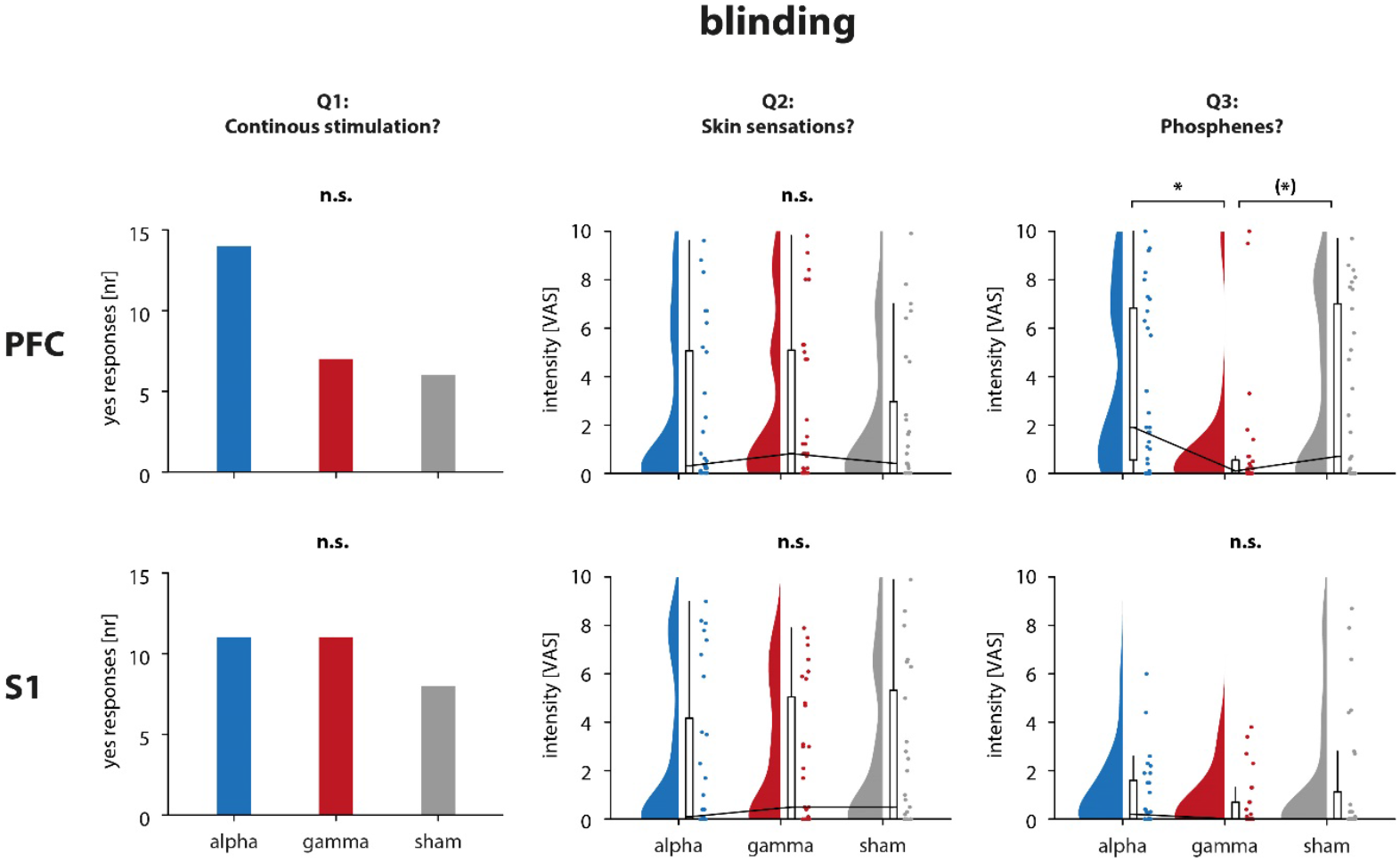
Blinding. Results of blinding questionnaires assessing the perception of a continuous stimulation (Q1), the intensity of skin sensations (Q2), and the intensity of phosphenes (Q3) are displayed for PFC and S1 locations. Phosphenes were stronger for tACS over PFC than over S1 (N = 29; p = .007; Friedman test; not shown here). In addition, phosphenes differed between alpha, gamma, and sham stimulation for tACS over PFC but not over S1 (N = 29; PFC: p = .012; S1: p = .910; Friedman tests). Post hoc tests showed that phosphenes were significantly stronger in the PFC alpha condition than in the PFC gamma condition (N = 29; p = .006; Wilcoxon matched-pairs signed rank tests; Bonferroni-corrected p-value). (*> .05 < p < .10, * p < .05; n.s., not significant; PFC, prefrontal cortex; S1, somatosensory cortex; tACS, transcranial alternating current stimulation; Q1-3, question 1-3; VAS, visual analogue scale.

**Supplementary Fig. 2.**
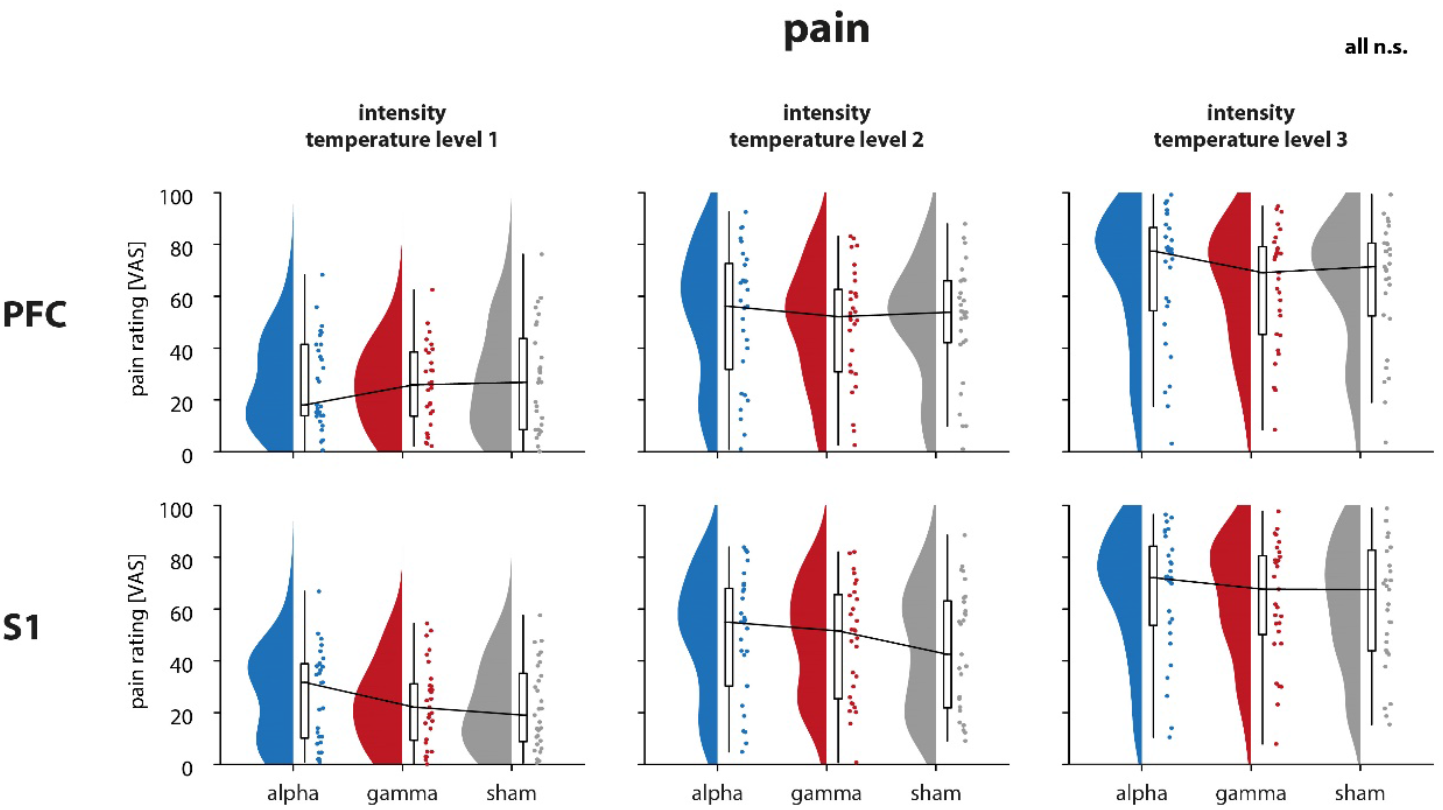
tACS effects on pain per temperature level. tACS effects on averaged pain ratings (0-100; VAS) at each temperature level (low, medium, high) are shown. Raincloud plots [2] show un-mirrored violin plots displaying the probability density function of the data, boxplots, and individual data points. Boxplots depict the sample median as well as first (Q1) and third quartiles (Q3). Whiskers extend from Q1 to the smallest value within Q1 – 1.5* interquartile range (IQR) and from Q3 to the largest values within Q3 + 1.5* IQR. None of the analyses revealed significant differences between alpha, gamma, and sham stimulation indicating no tACS effects on the perceived pain intensity at specific temperature levels (N = 29; PFC_leve1_: p = .857, PF_Clevel2_: p = .857, PF_Clevel3_: p = 1.0, S1_level2_: p = .857, S1_level3_: p = .864; Friedman tests; FDR-corrected p-values). n.s., not significant; PFC, prefrontal cortex; S1, somatosensory cortex; tACS, transcranial alternating current stimulation; VAS, visual analogue scale.

**Supplementary Table 1.** Statistical results of pain and autonomic activity analyses (uncorrected p-values).

|  | main analysis |  |  |  | responder analysis |  |  |  |  |  |  |  | peak freq analysis |  |
| --- | --- | --- | --- | --- | --- | --- | --- | --- | --- | --- | --- | --- | --- | --- |
|  |  |  |  |  | alpha responders |  |  |  | gamma responders |  |  |  |  |  |
|  | PFC |  | S1 |  | PFC |  | S1 |  | PFC |  | S1 |  | S1 |  |
|  | Z(2) | p | Z(2) | p | Z(2) | p | Z(2) | p | Z(2) | p | Z(2) | p | Z(2) | p |
| **pain** |  |  |  |  |  |  |  |  |  |  |  |  |  |  |
| **intensity** |  |  |  |  |  |  |  |  |  |  |  |  |  |  |
| 8 min | 0.90 | .639 | 1.10 | .576 | -0.68 | .496 | -1.07 | .286 | -1.33 | .184 | -0.38 | .701 | -1.22 | .221 |
| time-resolved | - | no cl. | - | .319 | - | .130 | - | .026 | - | .283 | - | .329 | - | .070 |
| temp level 1 | 1.93 | .381 | 0.21 | .902 | -0.57 | .570 | -0.59 | .557 | -1.20 | .231 | -0.11 | .917 | -0.45 | .650 |
| temp level 2 | 1.93 | .381 | 2.14 | .343 | -0.74 | .460 | -1.29 | .199 | -1.15 | .248 | -0.59 | .552 | -1.22 | .221 |
| temp level 3 | 0.07 | .966 | 1.31 | .519 | -0.06 | .955 | -0.89 | .372 | -0.76 | .446 | -0.45 | .650 | -1.22 | .221 |
| **variability** |  |  |  |  |  |  |  |  |  |  |  |  |  |  |
| 8 min | 1.10 | .576 | 4.21 | .122 | -0.51 | .609 | -0.76 | .446 | -0.07 | .948 | -0.59 | .552 | -0.18 | .861 |
| time-resolved | - | .254 | - | .454 | - | .136 | - | no cl. | - | no cl. | - | .162 | - | no cl. |
| **relation to thermal stimulation** |  |  |  |  |  |  |  |  |  |  |  |  |  |  |
| 8 min | 0.48 | .790 | .48 | .790 | -0.06 | .955 | -0.94 | .349 | -1.42 | .157 | -0.80 | .422 | -0.94 | .345 |
| time-resolved | - | .106 | - | .337 | - | .024 | - | .168 | - | .099 | - | .193 | - | .252 |
| **autonomic activity** |  |  |  |  |  |  |  |  |  |  |  |  |  |  |
| **nSF** |  |  |  |  |  |  |  |  |  |  |  |  |  |  |
| 8 min | 3.62 | .164 | 0.92 | .631 | -0.41 | .683 | -2.15 | .031 | -0.78 | .435 | -0.11 | .916 | -0.94 | .347 |
| time-resolved | - | no cl. | - | no cl. | - | no cl. | - | .072 | - | no cl. | - | no cl. | - | .088 |
| **HR** |  |  |  |  |  |  |  |  |  |  |  |  |  |  |
| 8 min | 1.31 | .519 | 2.48 | .289 | -0.85 | .394 | -0.54 | .586 | -1.11 | .267 | -0.25 | .807 | -0.66 | .507 |
| time-resolved | - | no cl. | - | no cl. | - | no cl. | - | no cl. | - | no cl. | - | no cl. | - | no cl. |

*Note.* Results of Friedmann tests and cluster-based permutation statistics. Cluster-based permutation statistics were used for time-resolved analyses only. Uncorrected p-values are shown. HR, heart rate; no cl., no cluster found; nSF, number of skin conductance fluctuations; peak freq analysis, peak frequency-informed analysis; PFC, prefrontal cortex; S1, primary somatosensory cortex.

